## Supplementary figures and images for "Limited role of generation time changes in driving the evolution of mutation spectrum in humans"

### Fig1 Figure supplement 1

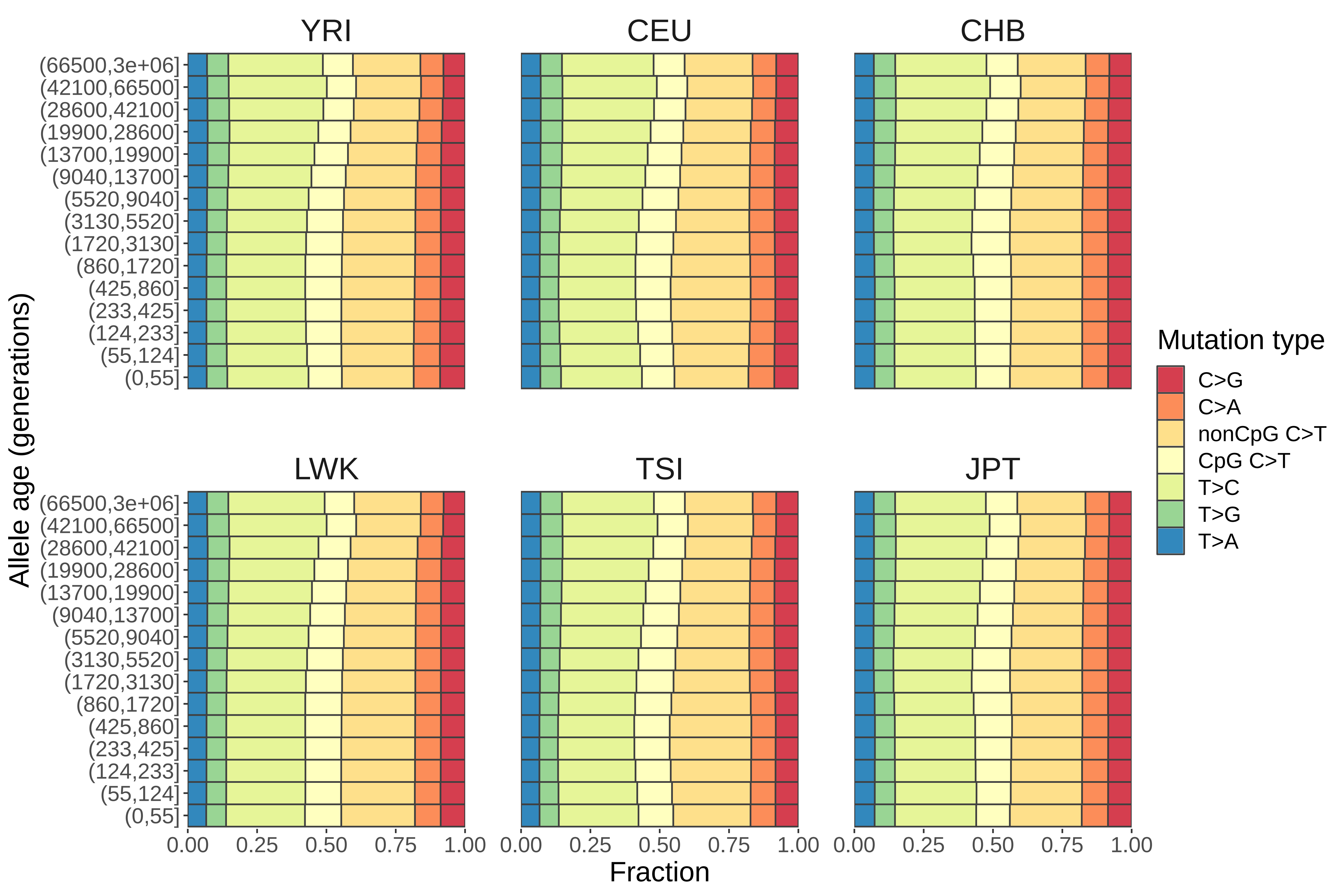

### Fig1 Figure supplement 2

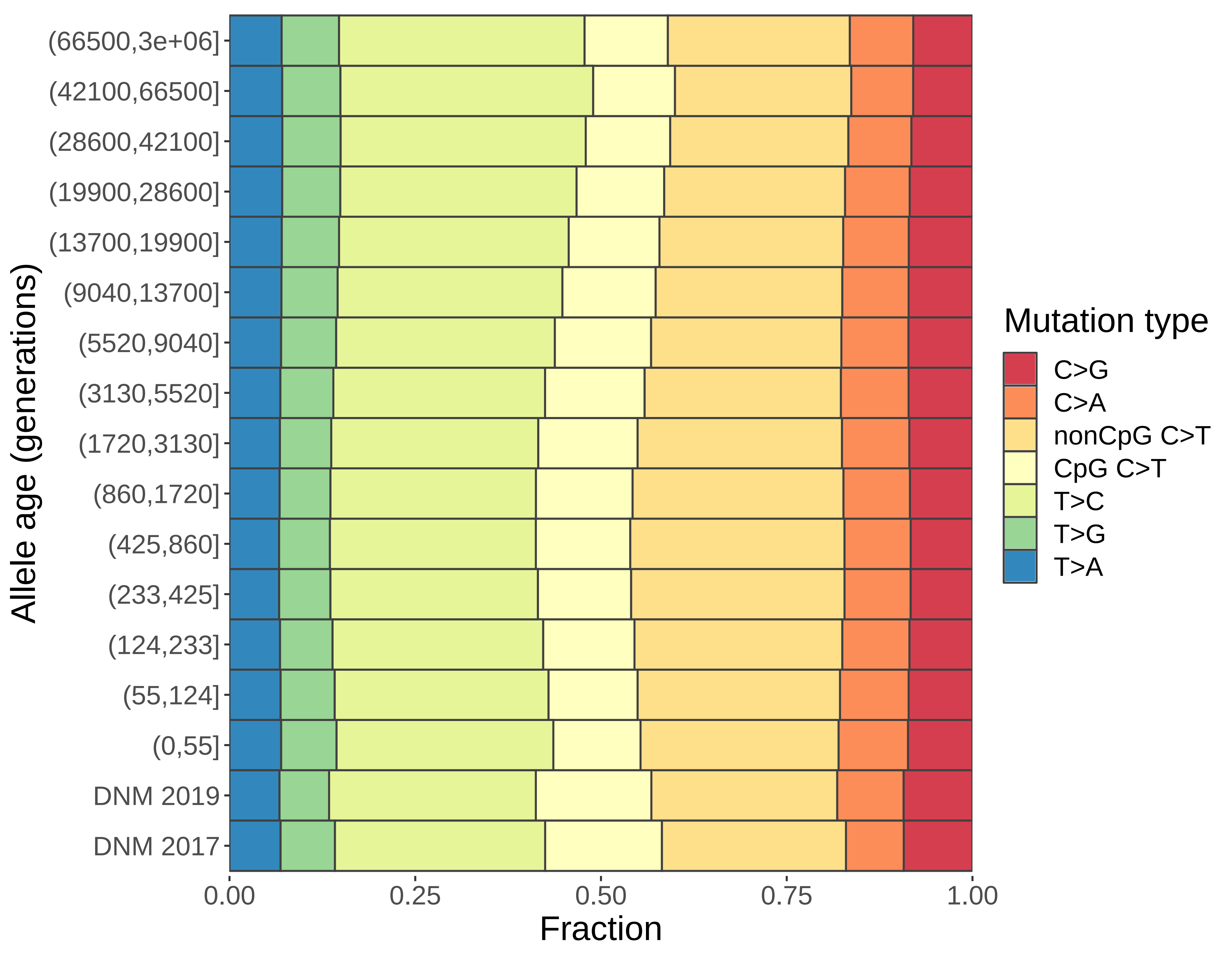

### Fig1 Figure supplement 3

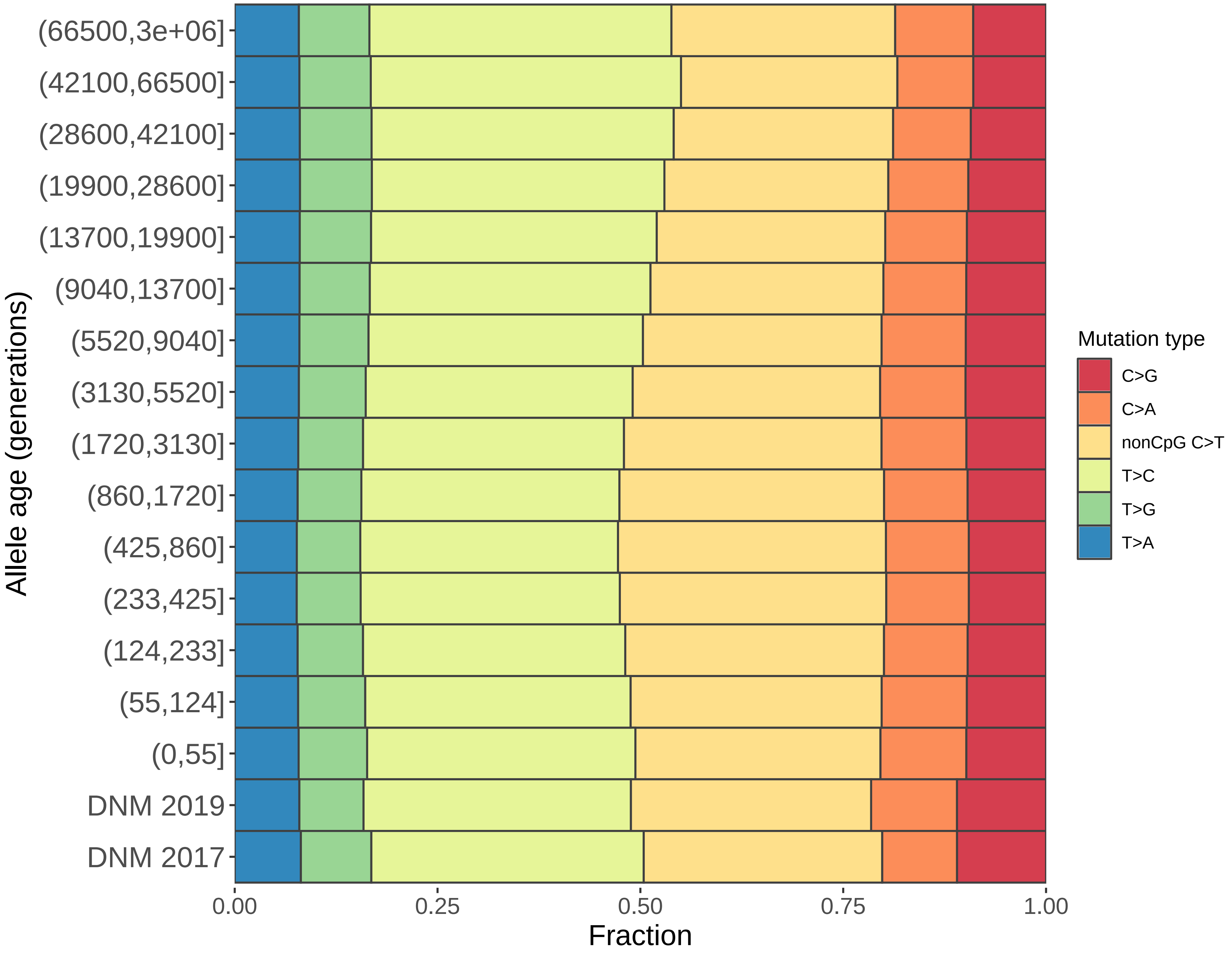

### Fig1 Figure supplement 4

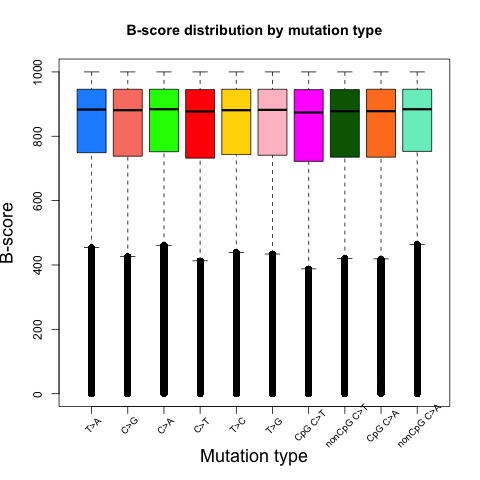

### Fig1 Figure supplement 5

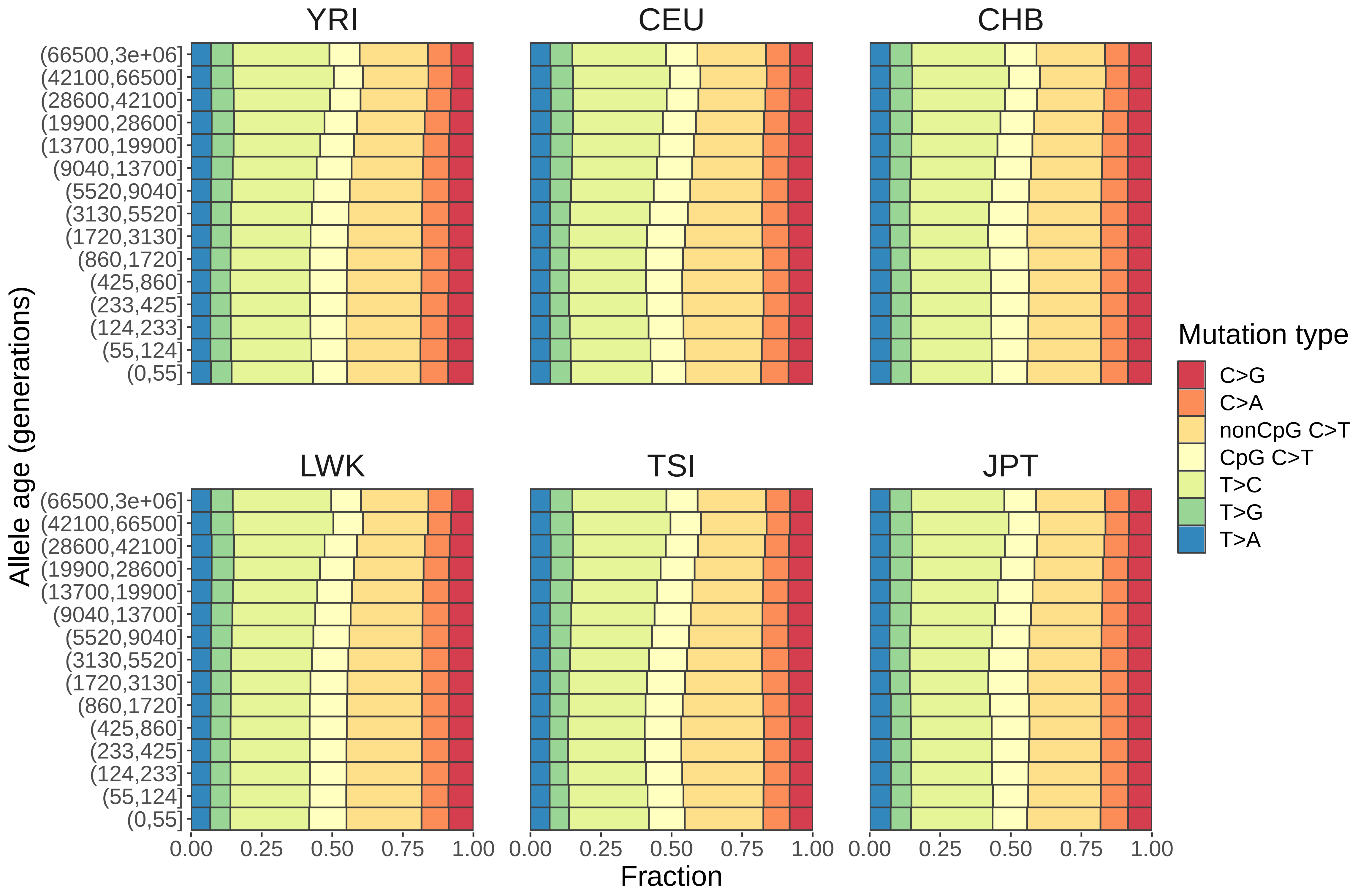

### Fig1 Figure supplement 6

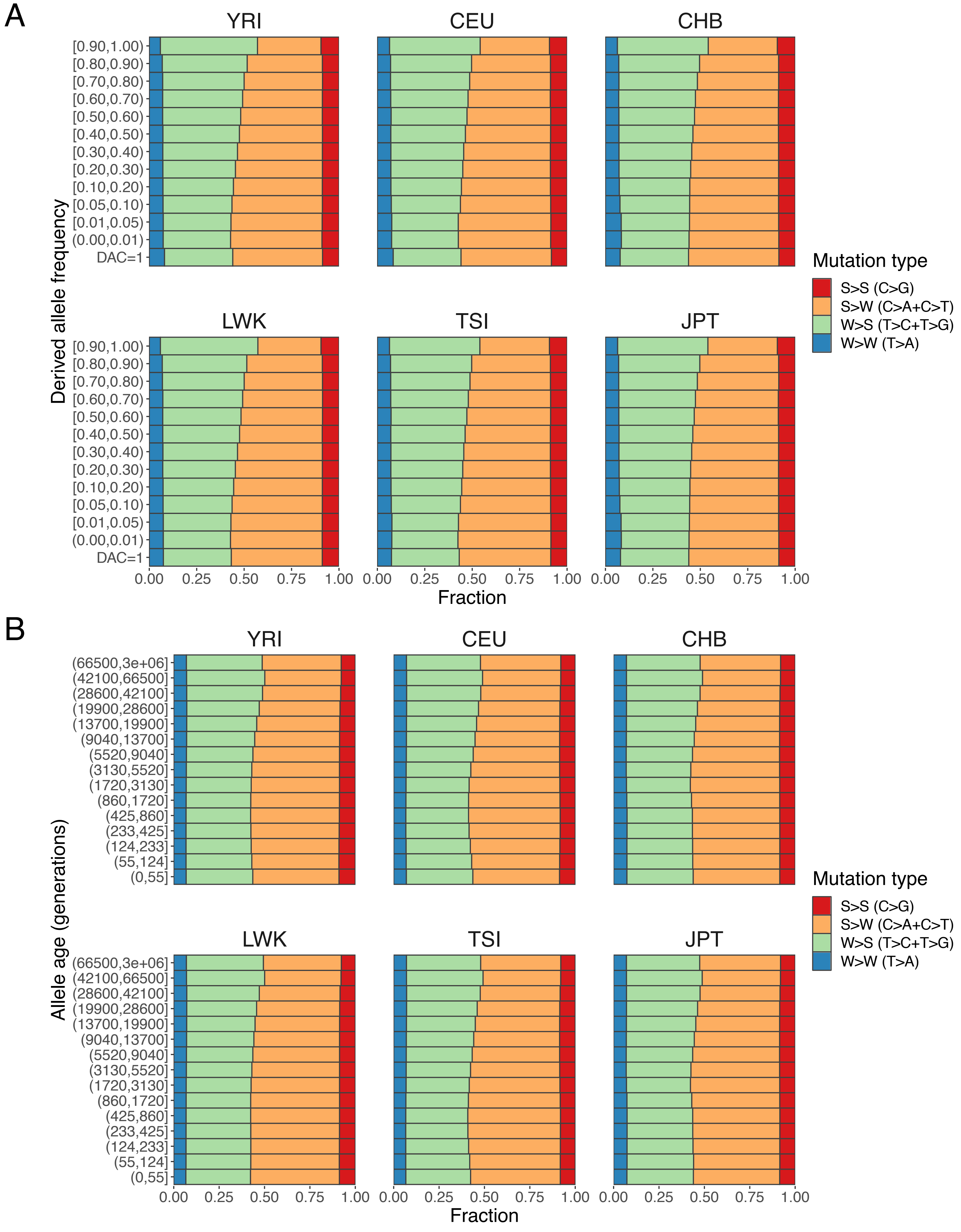

### Fig1 Figure supplement 7

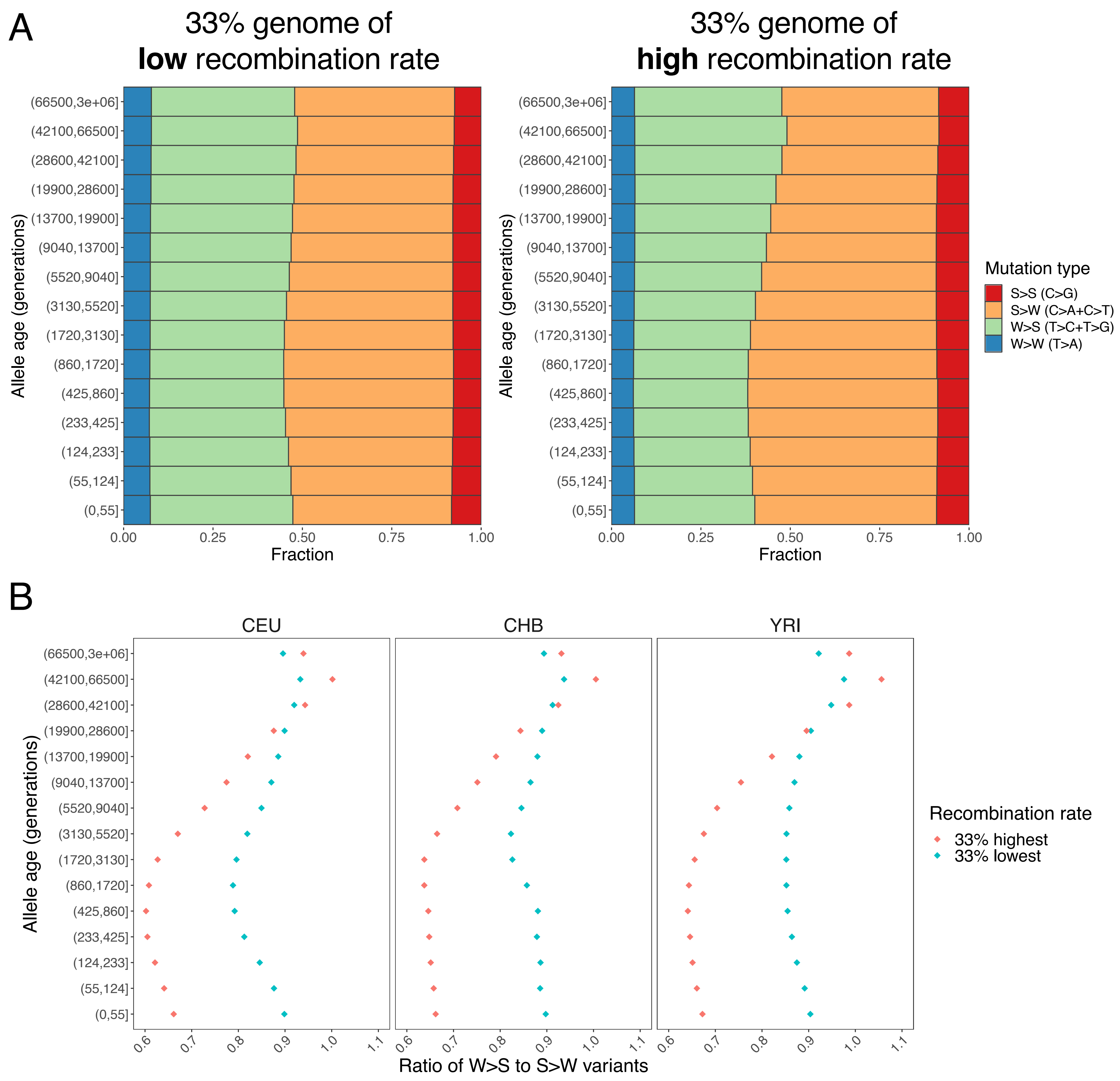

### Fig1 Figure supplement 8

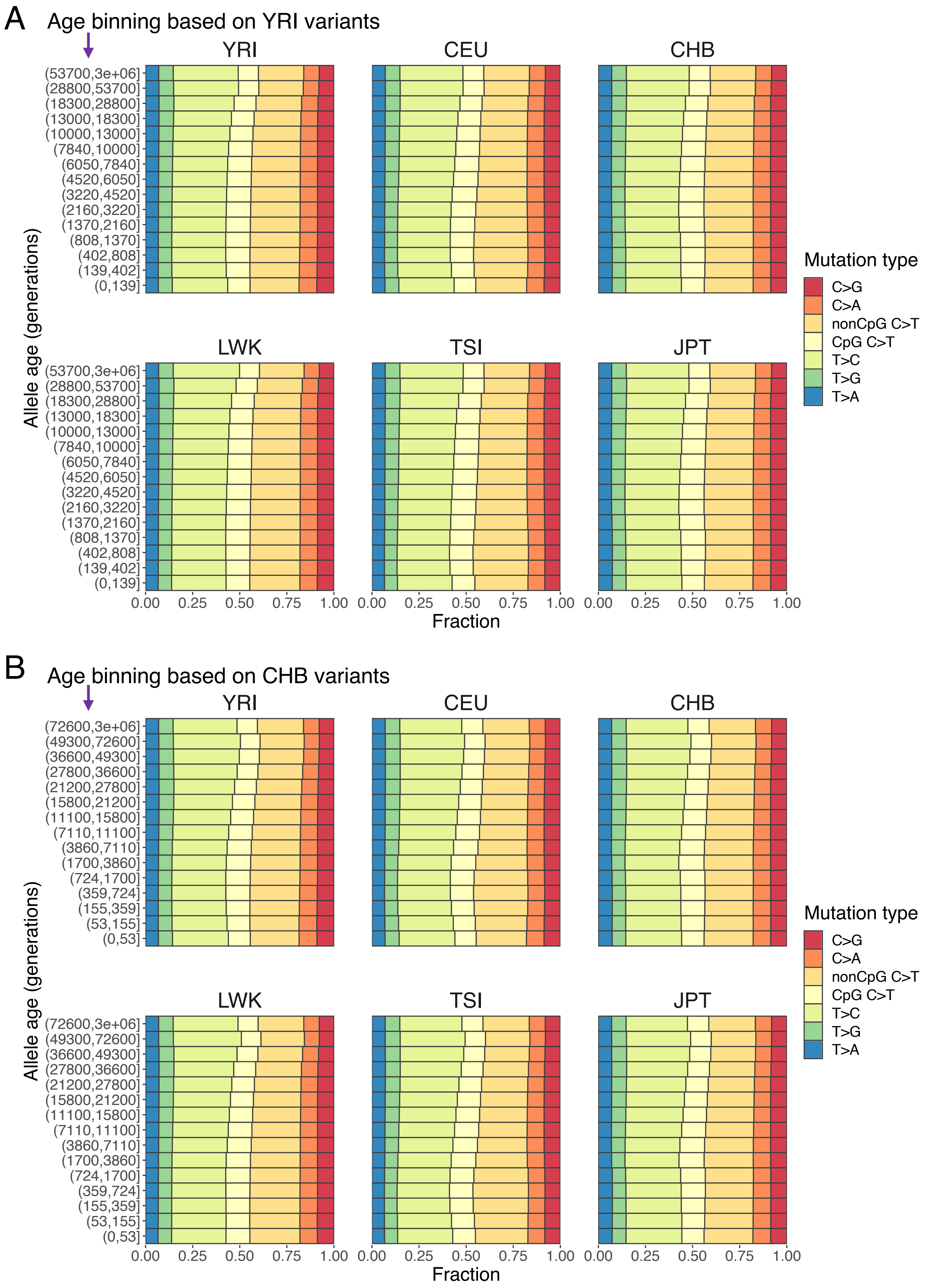

### Fig2 Figure supplement 1

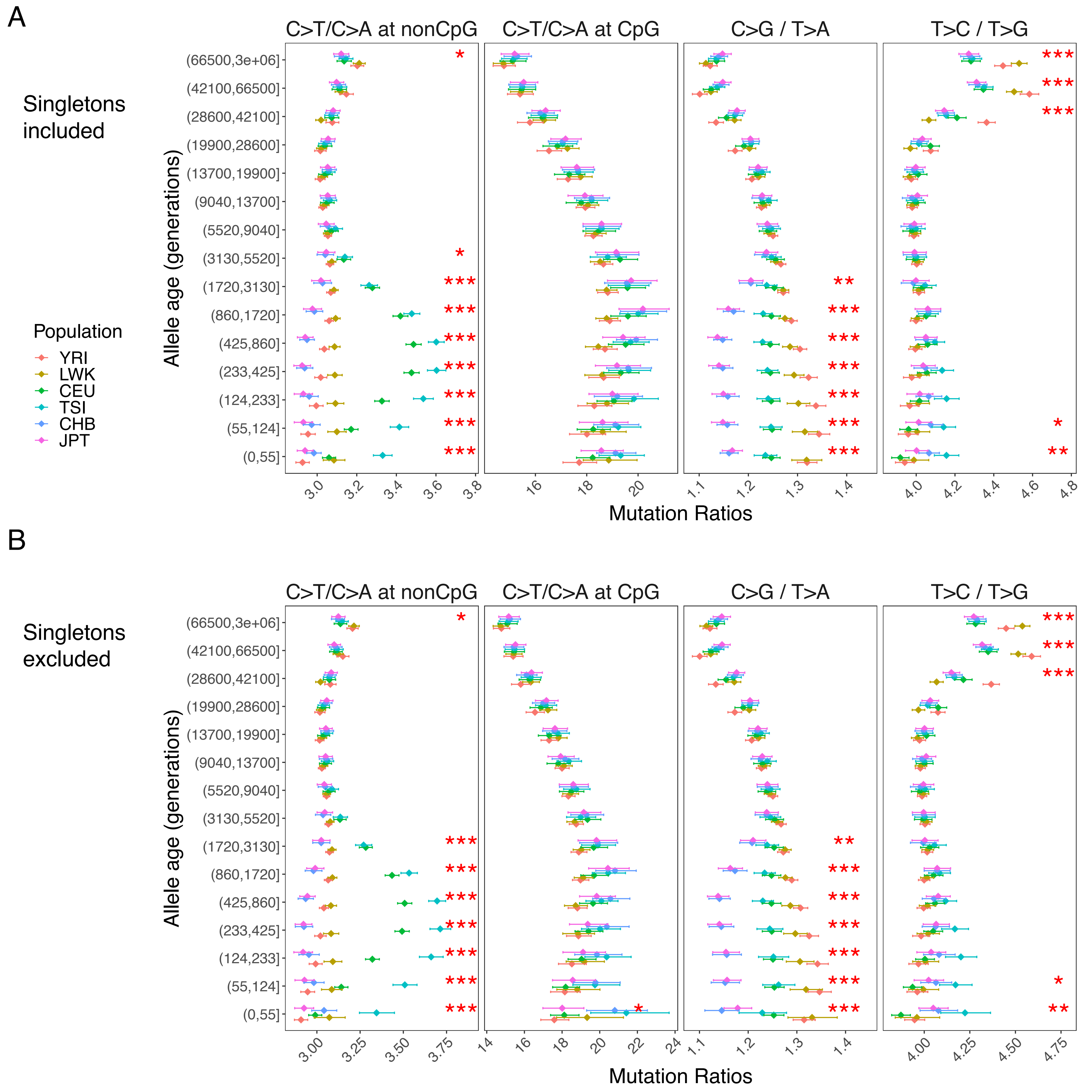

### Fig2 Figure supplement 2

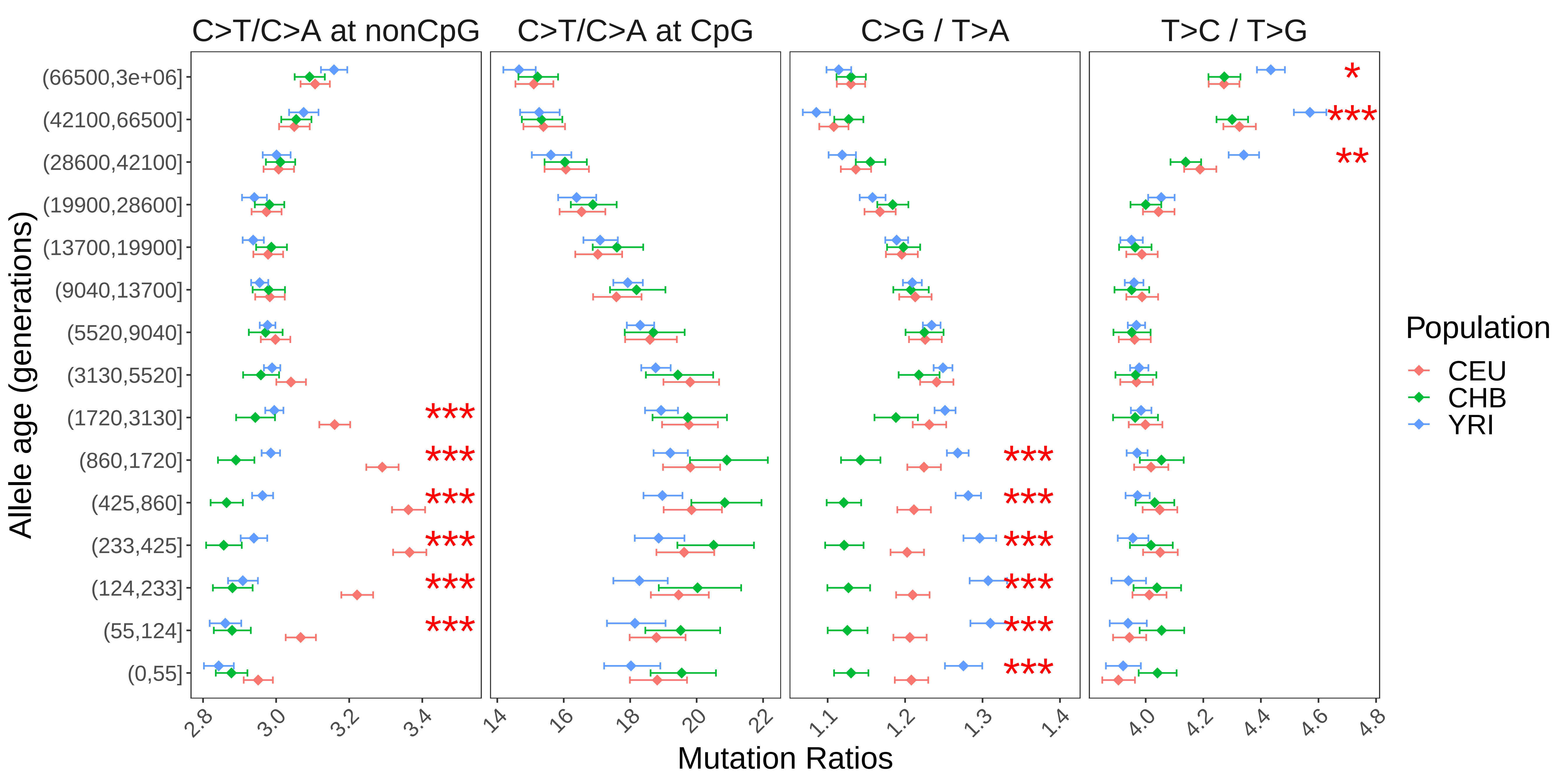

### Fig2 Figure supplement 3

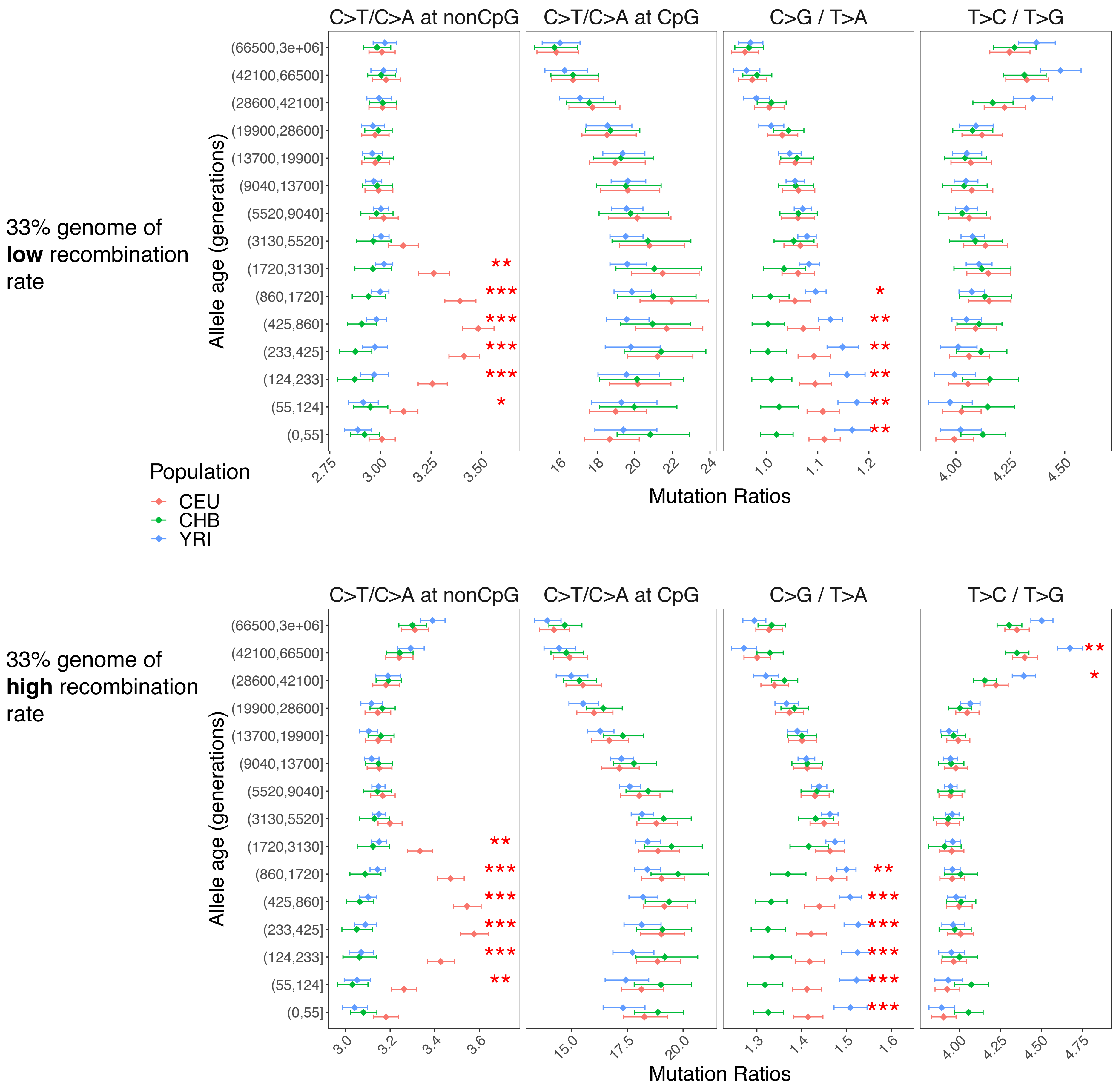

### Fig2 Figure supplement 4

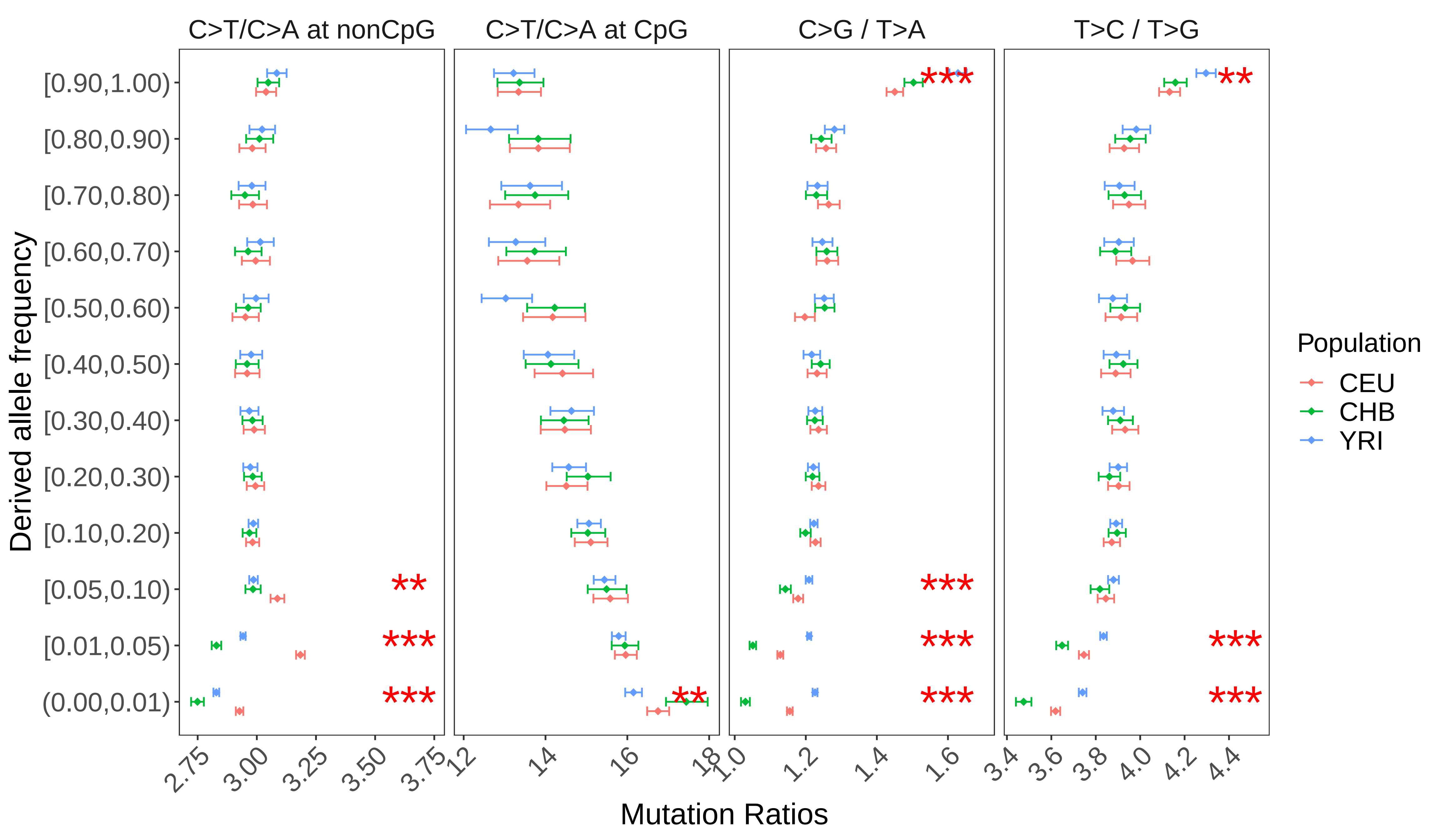

### Fig2 Figure supplement 5

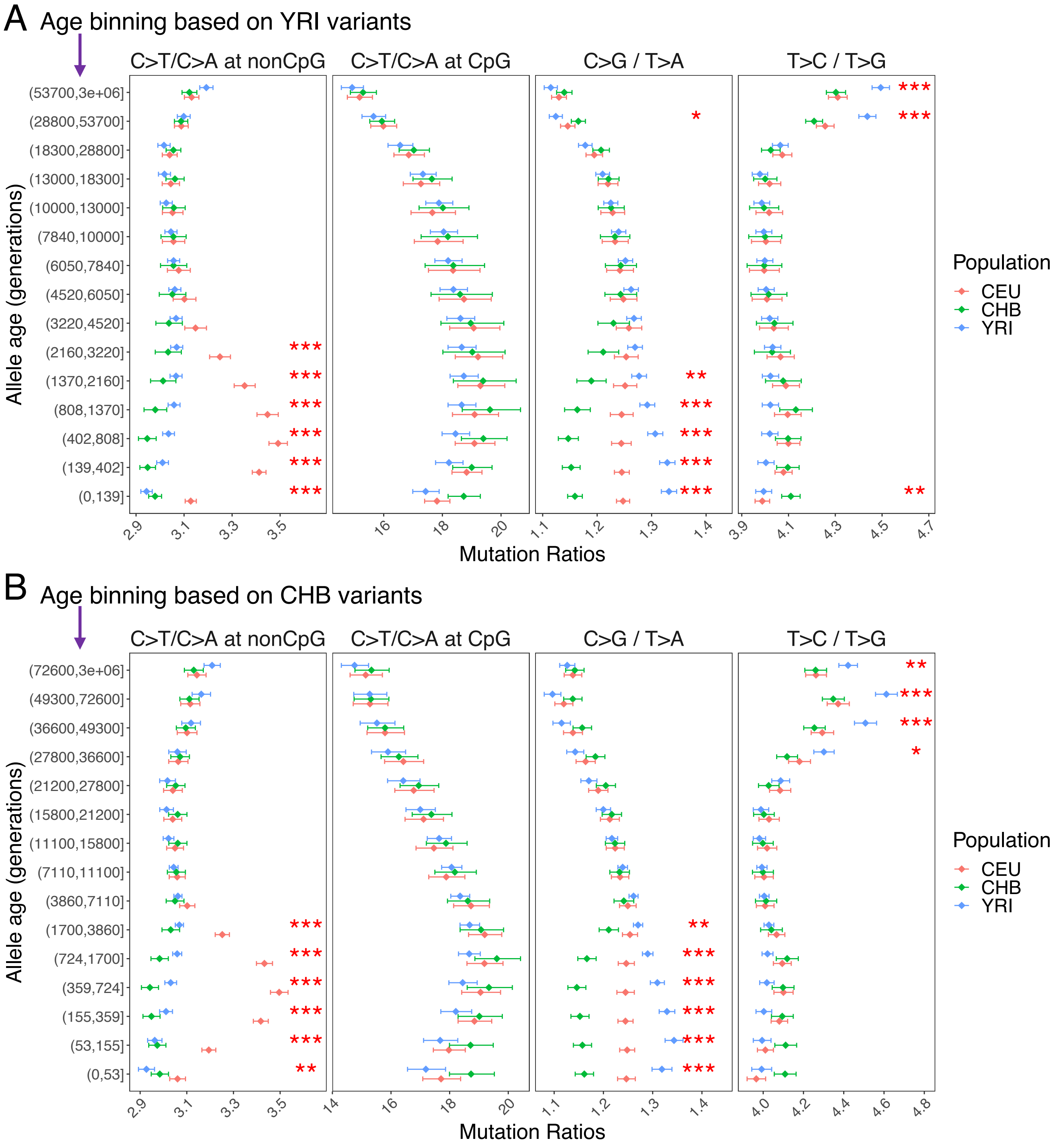

### Fig2 Figure supplement 6

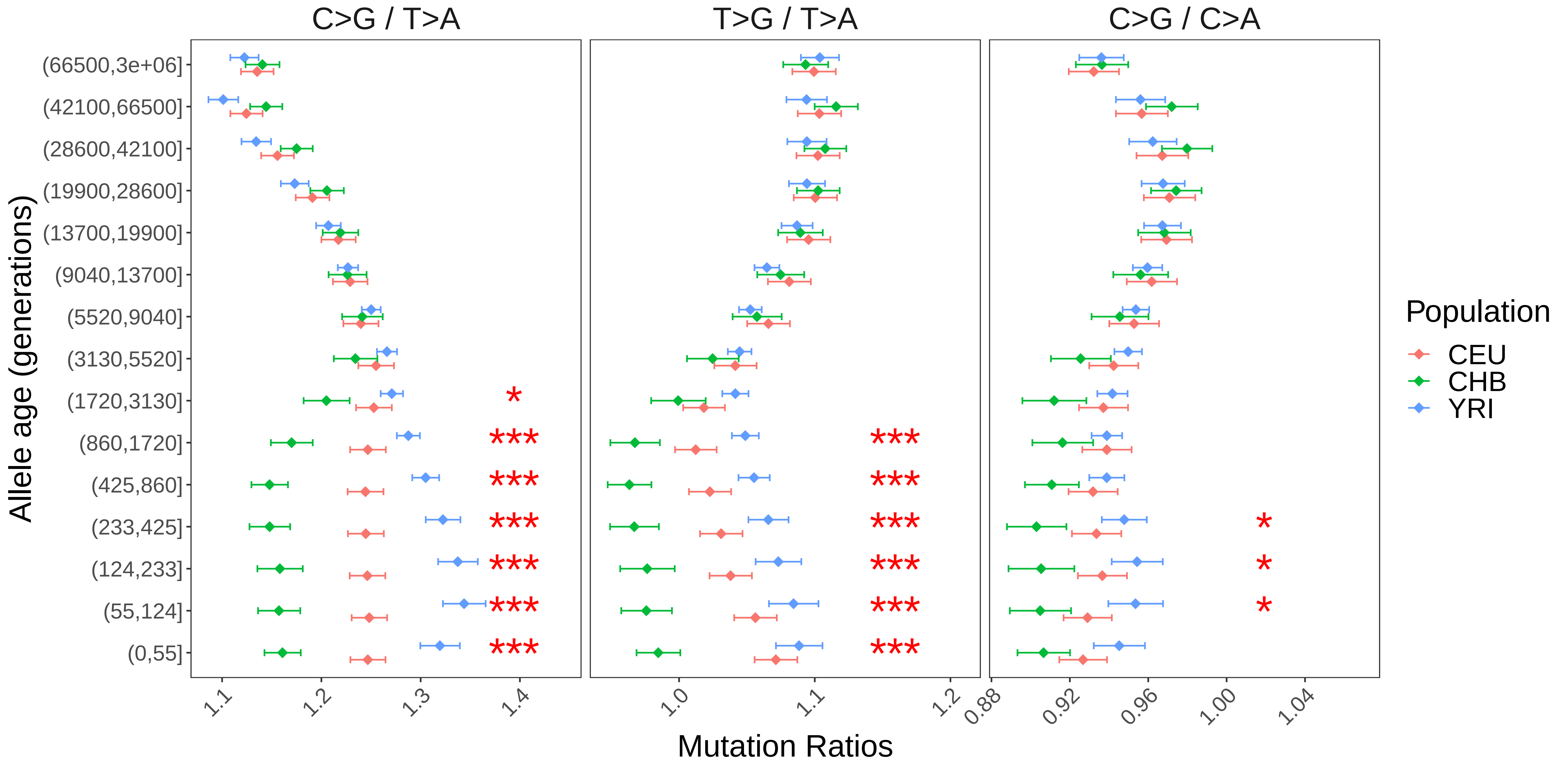

### Fig2 Figure supplement 7

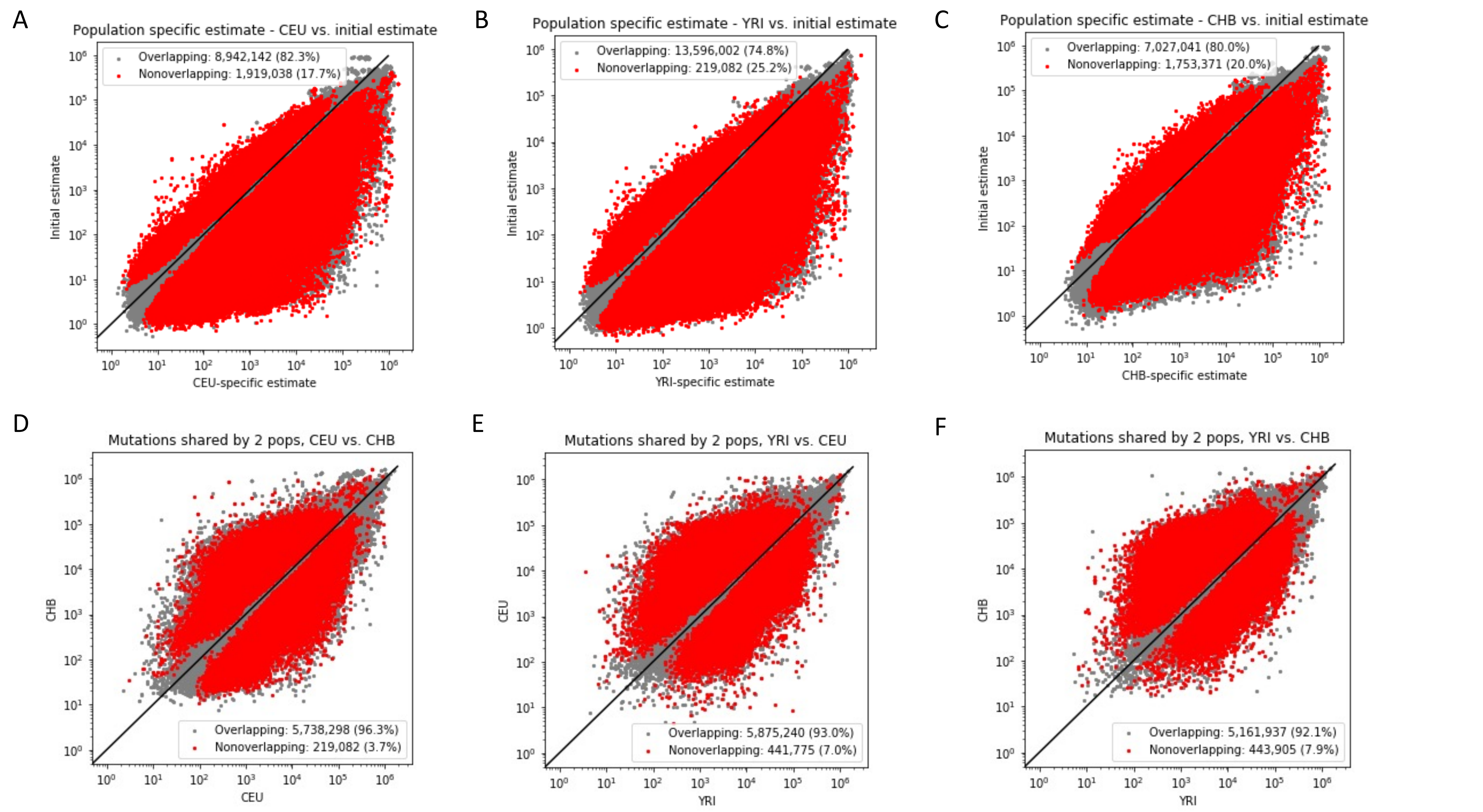

### Fig2 Figure supplement 8

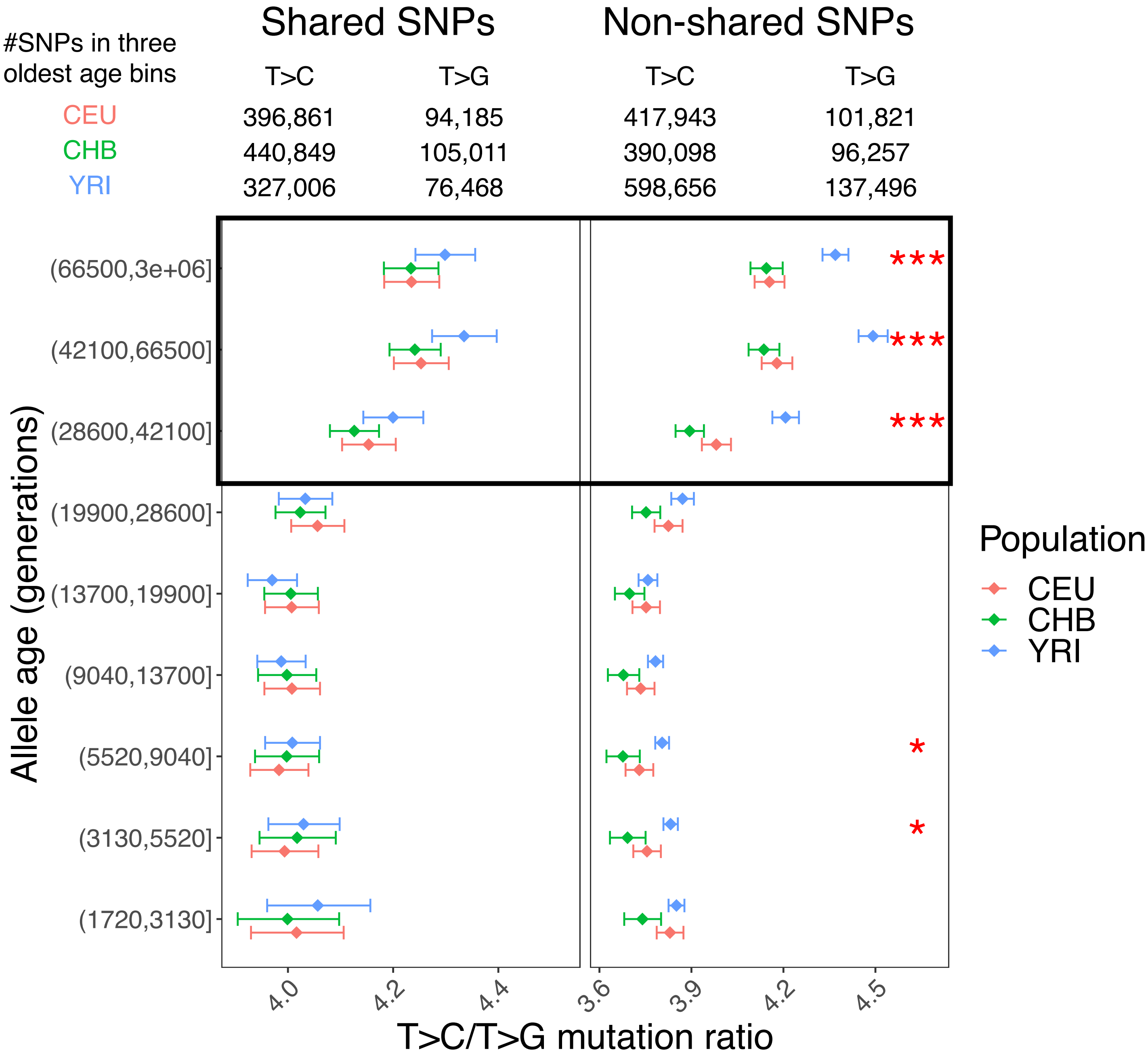

### Fig2 Figure supplement 9

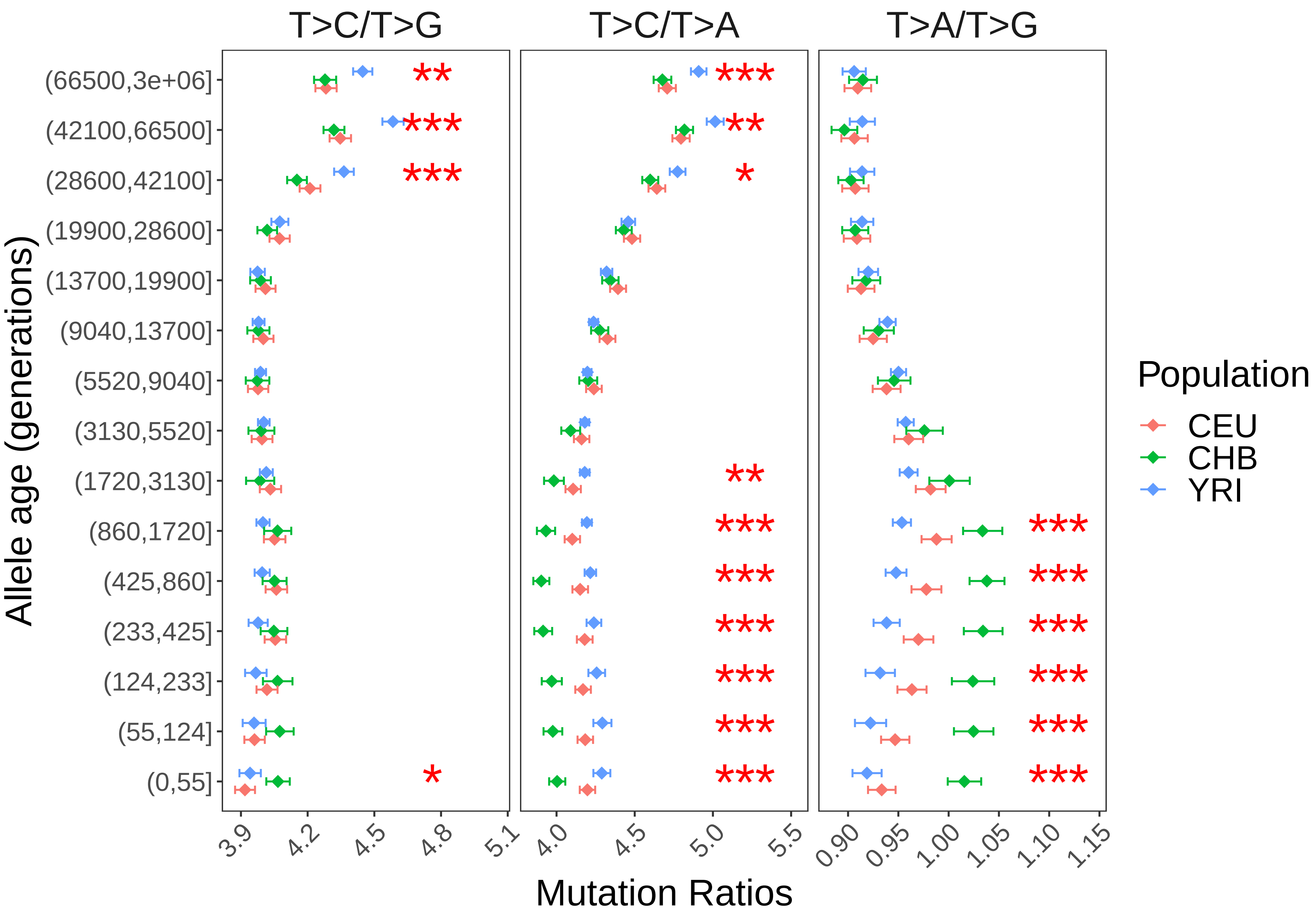

### Fig2 Figure supplement 10

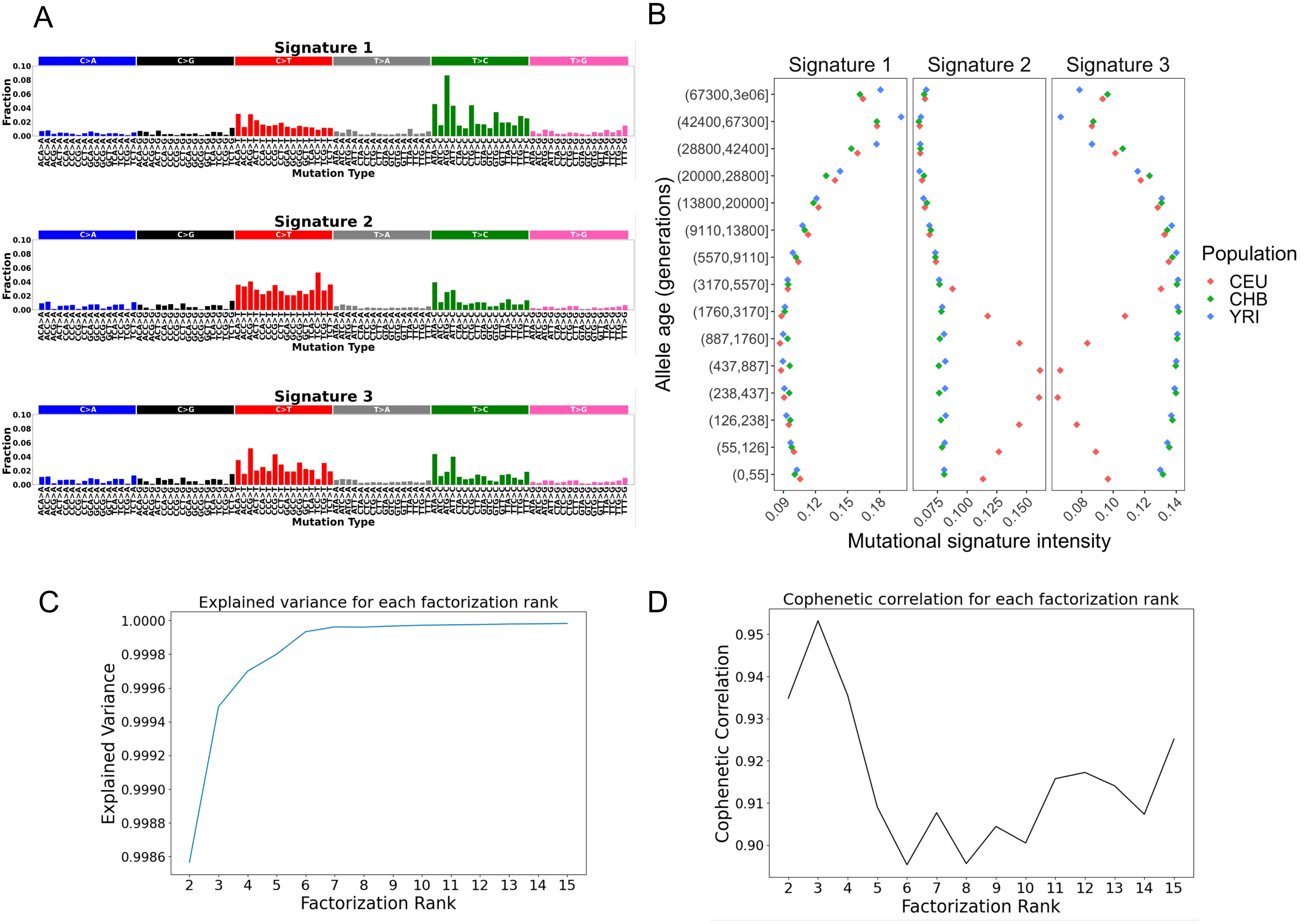

### Fig2 Figure supplement 11

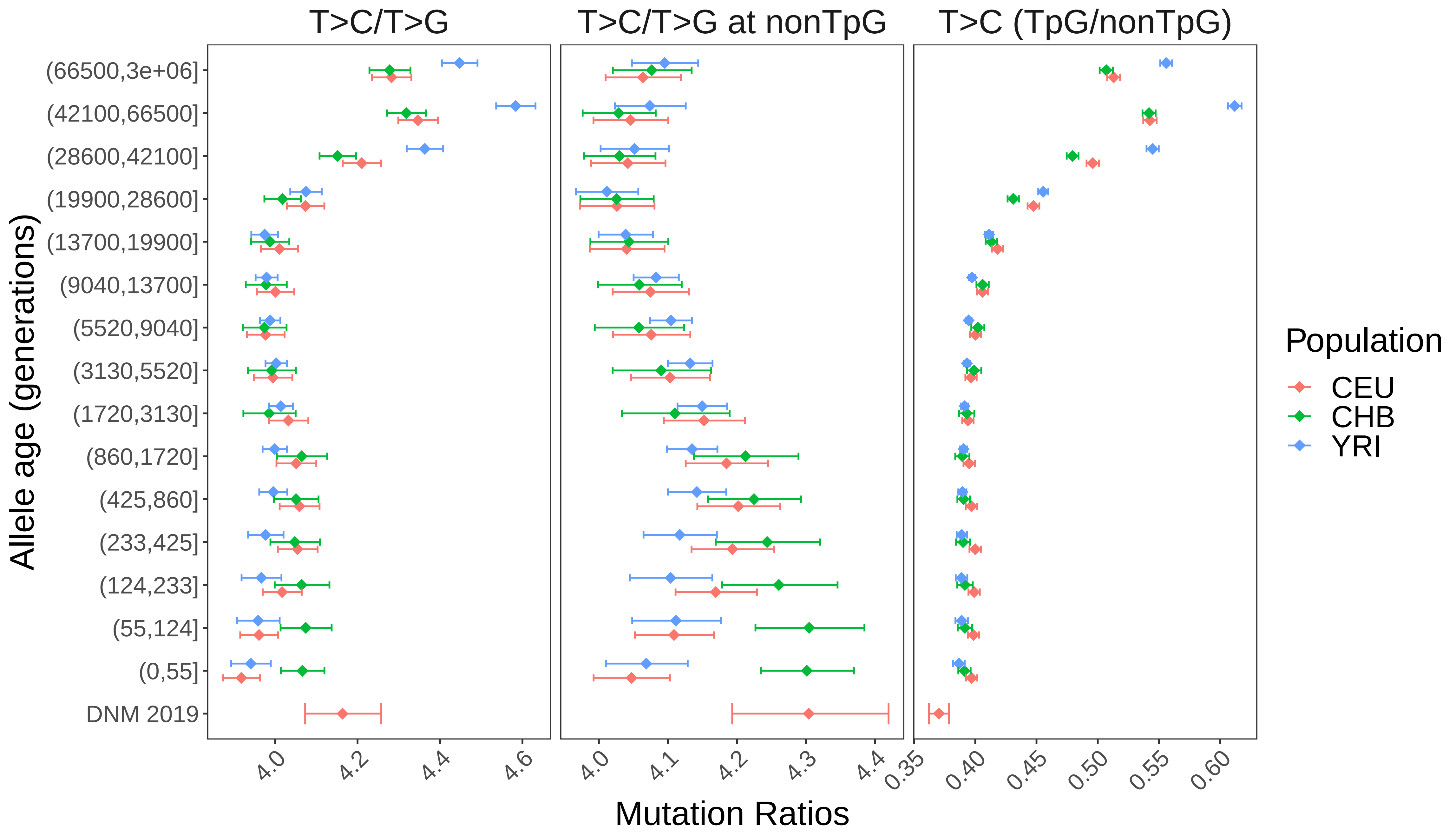

### Fig2 Figure supplement 12

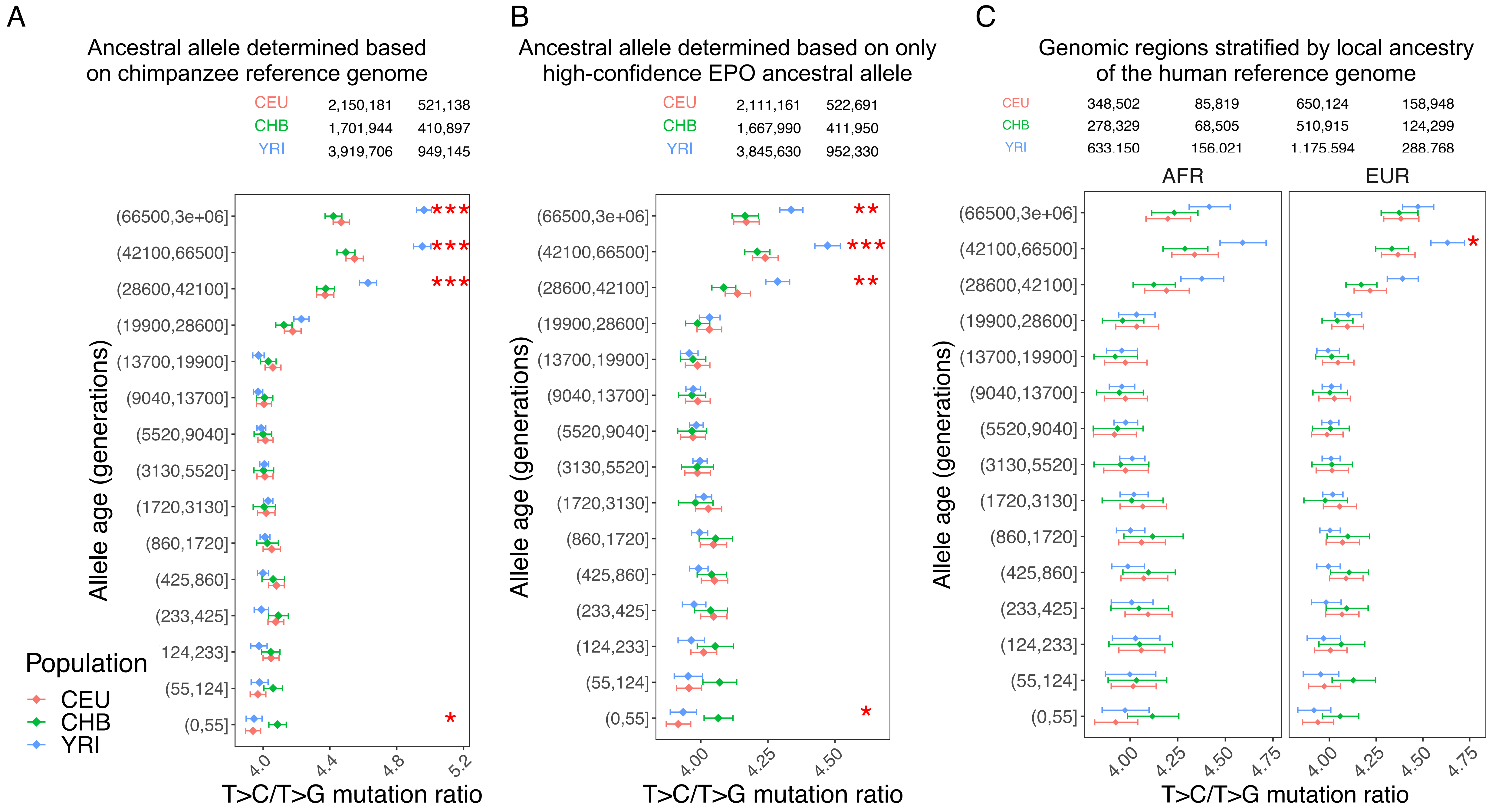

### Fig2 Figure supplement 13

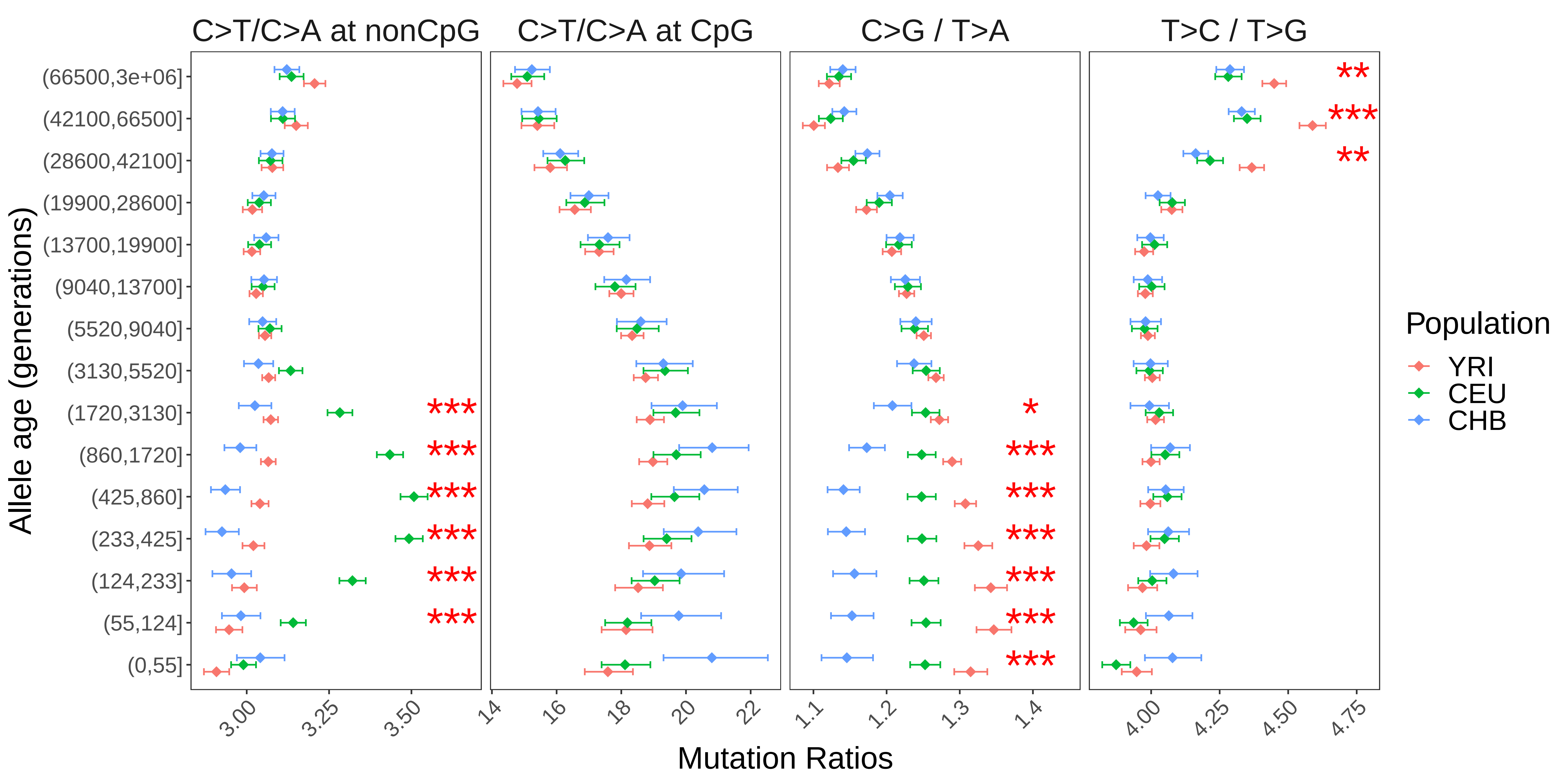

### Fig3 Figure supplement 1

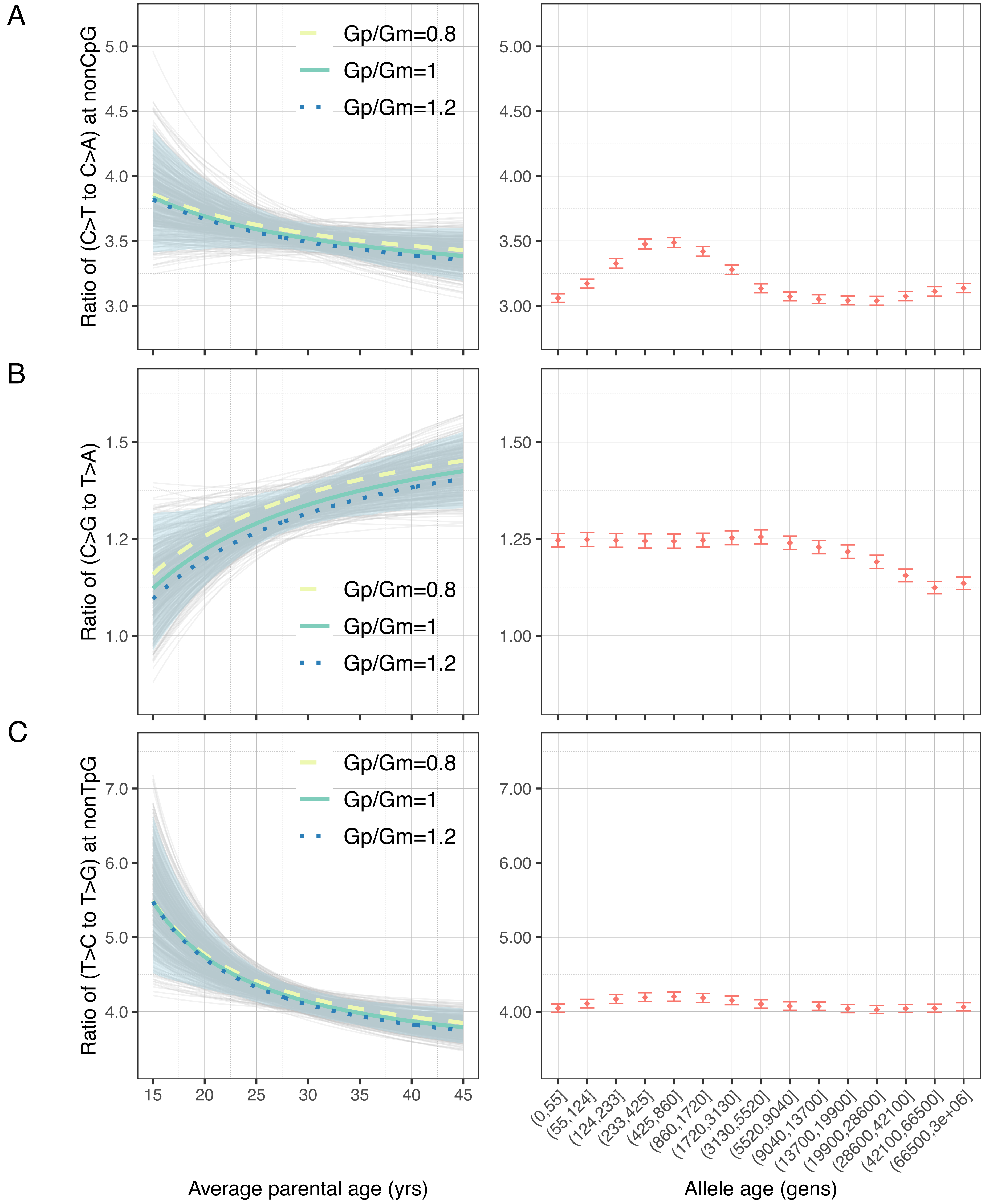

### Fig3 Figure supplement 2

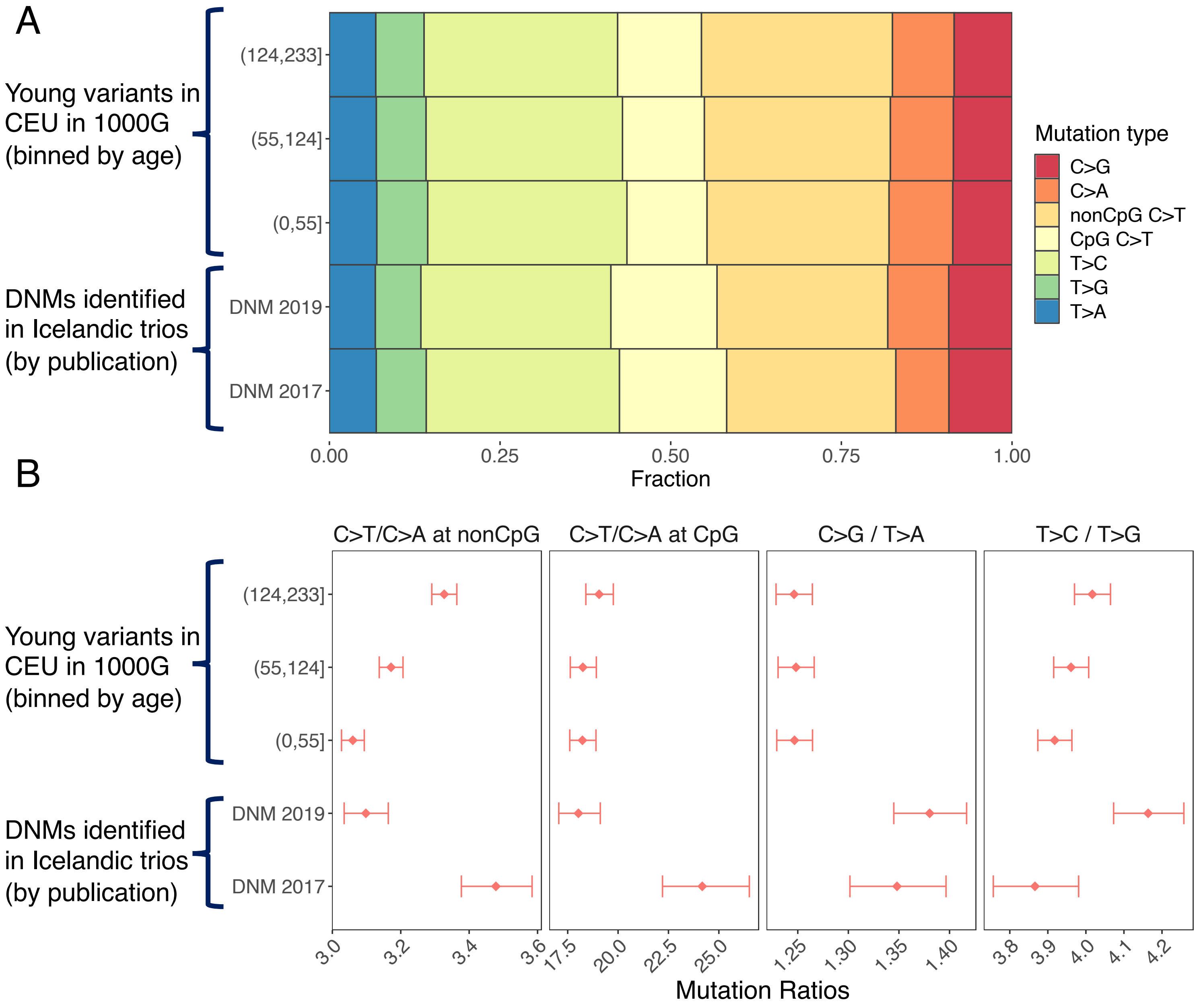

### Fig3 Figure supplement 3

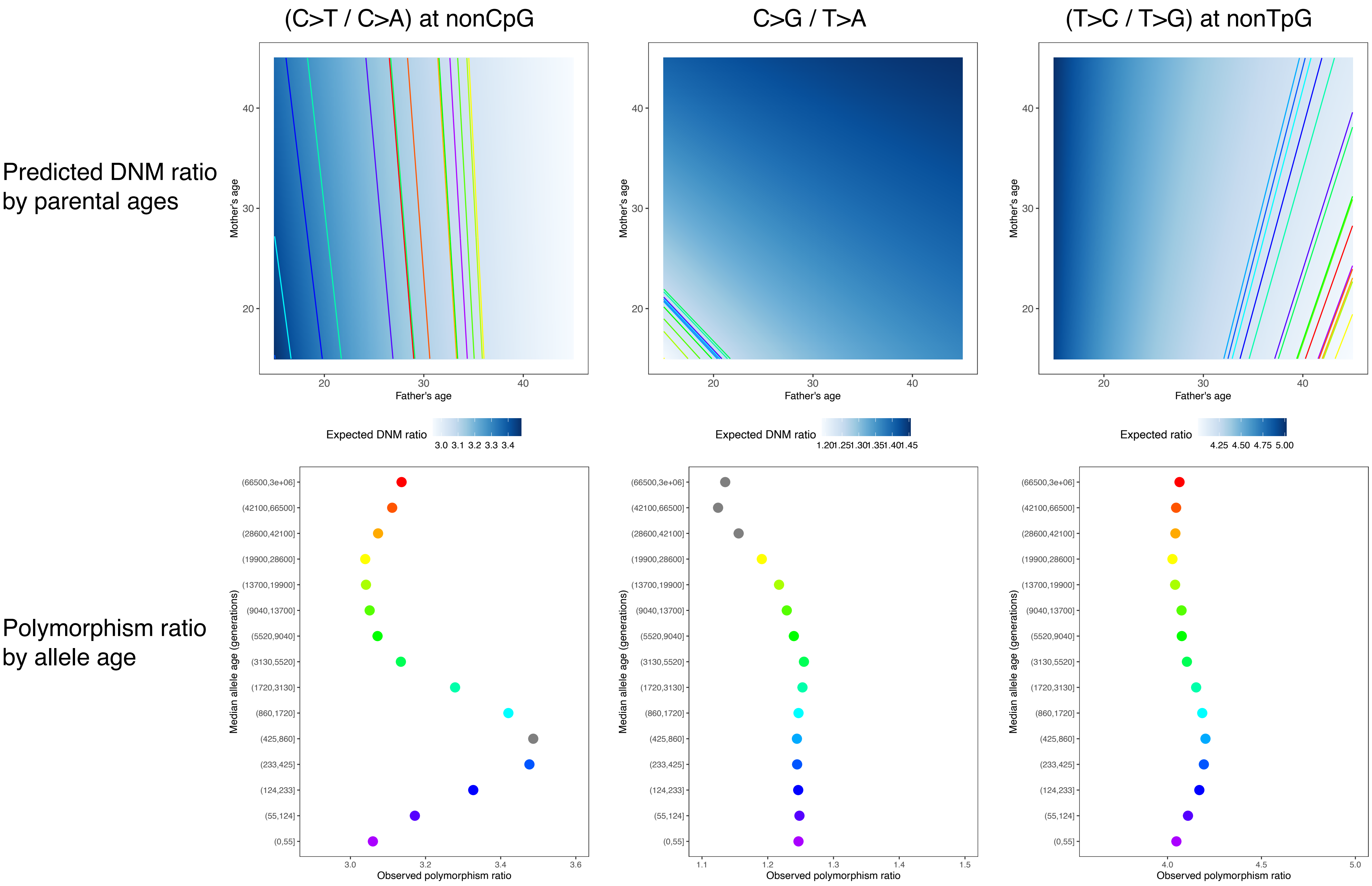

### Fig3 Figure supplement 4

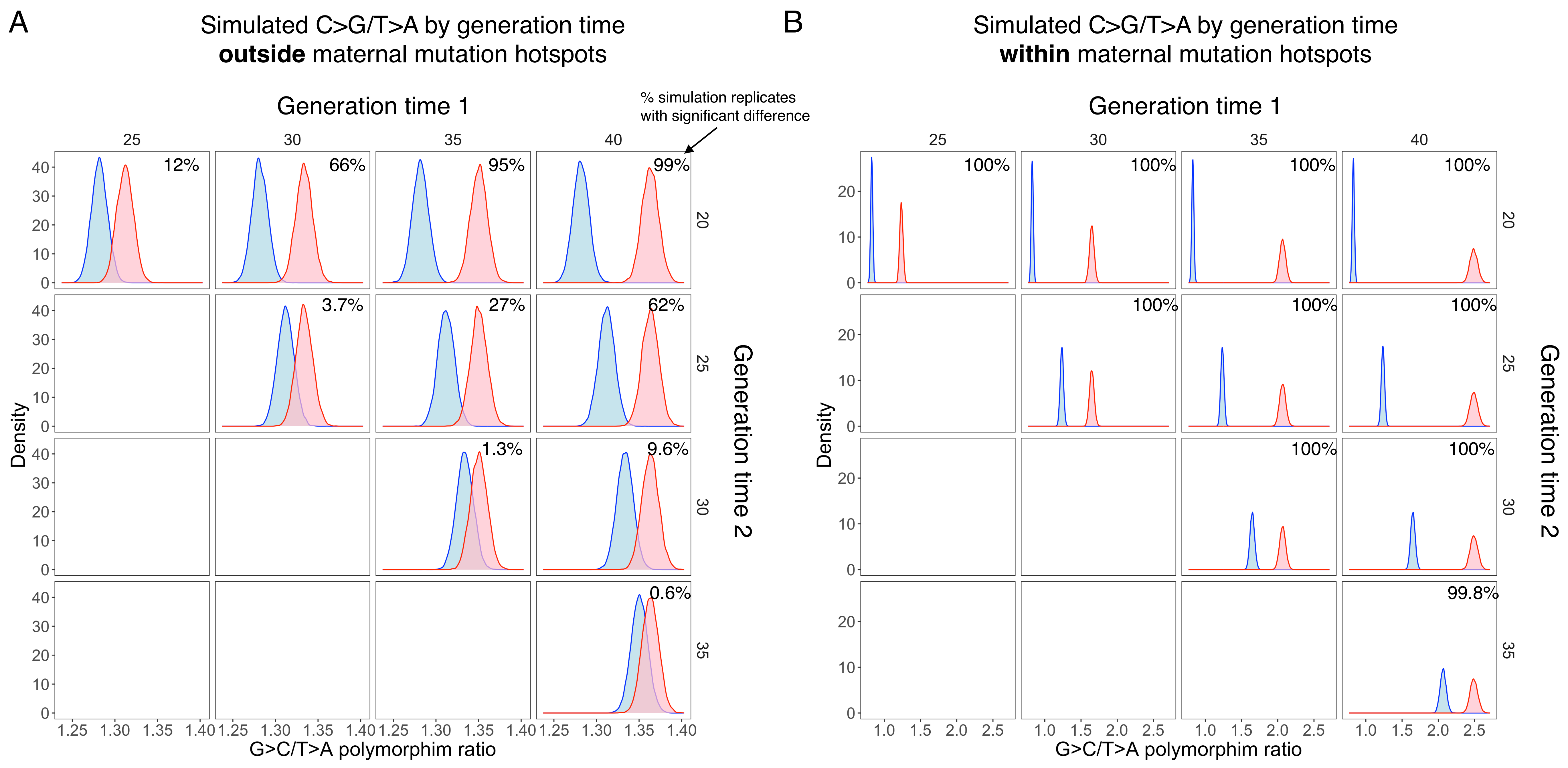

### Fig4 Figure supplement 1

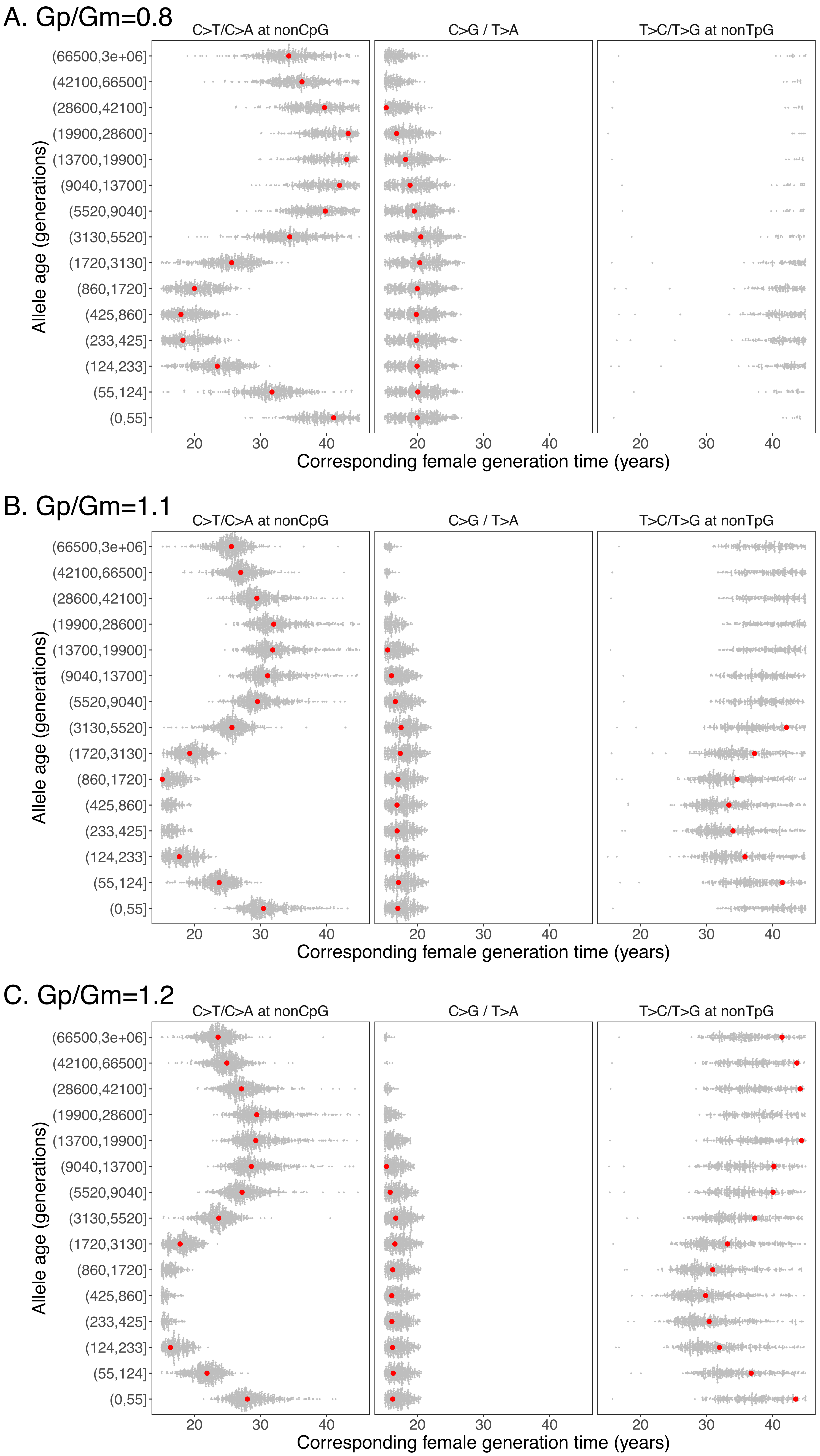

### Fig4 Figure supplement 2

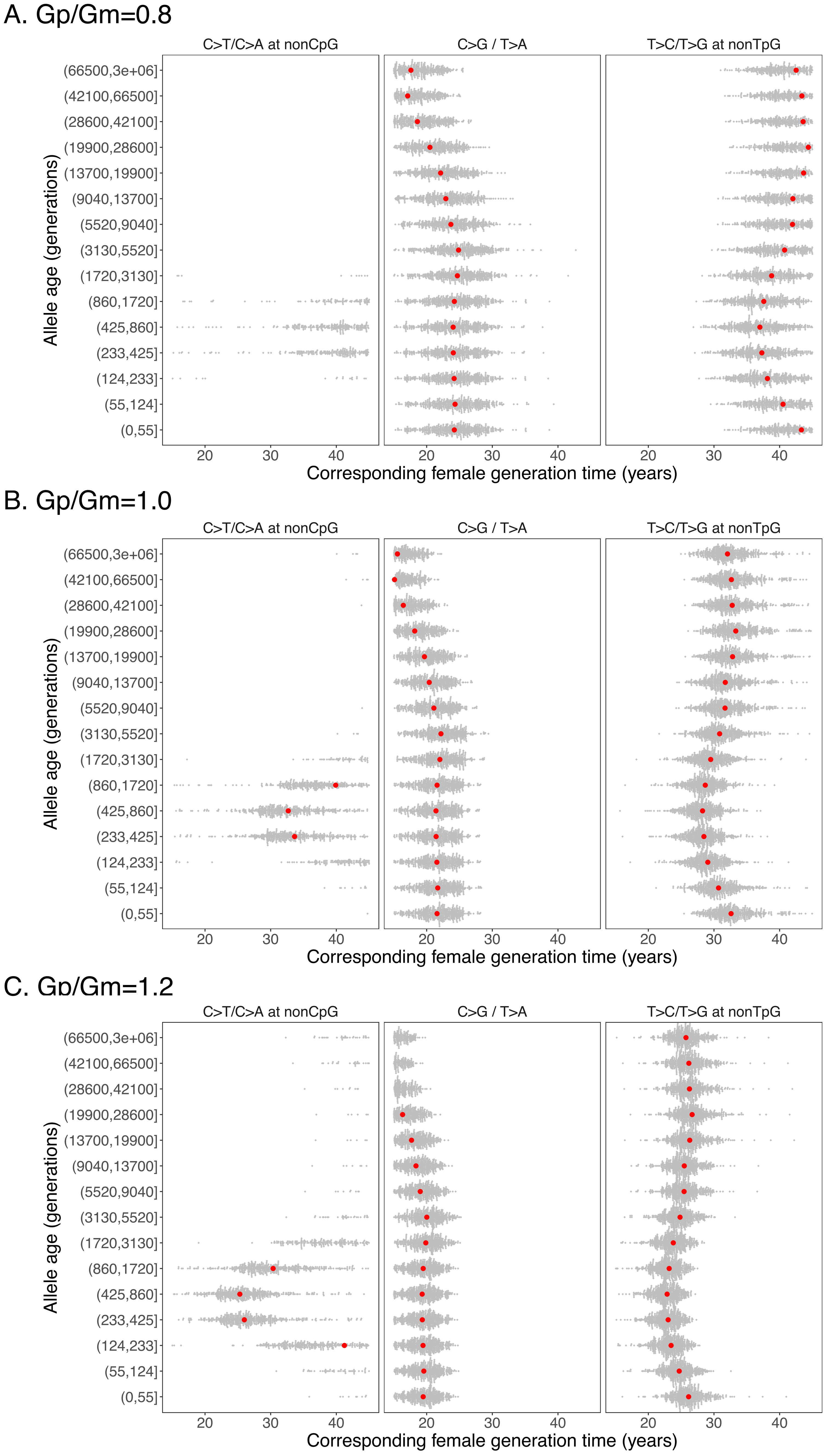

### Fig4 Figure supplement 3

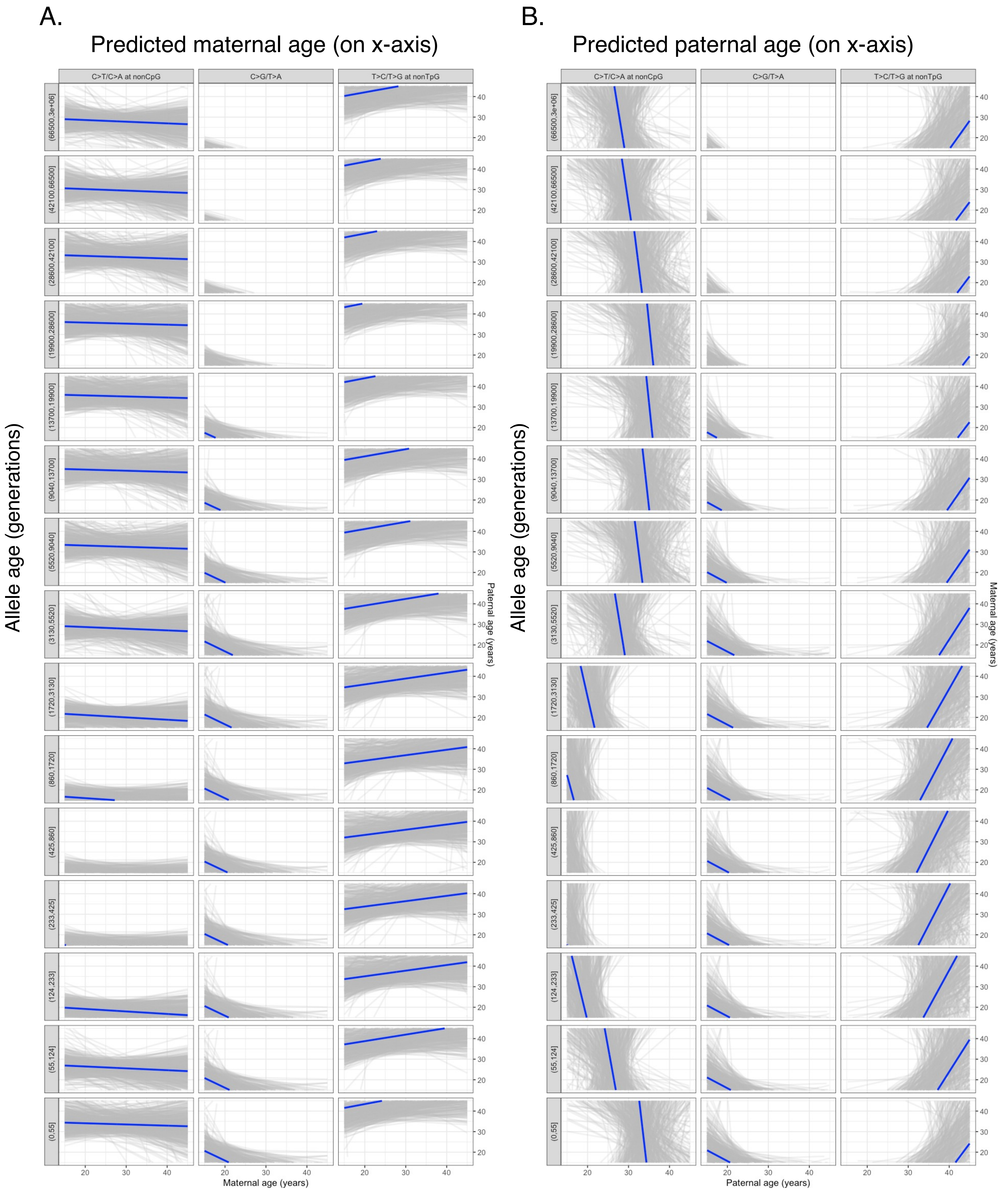
